# CD4^+^ T cells persist for years in the human small intestine and mediate robust T_H_1 immunity

**DOI:** 10.1101/863407

**Authors:** Raquel Bartolomé Casado, Ole J.B. Landsverk, Sudhir Kumar Chauhan, Frank Sætre, Kjersti Thorvaldsen Hagen, Sheraz Yaqub, Ole Øyen, Rune Horneland, Einar Martin Aandahl, Lars Aabakken, Espen S. Bækkevold, Frode L. Jahnsen

## Abstract

Studies in mice and humans have shown that CD8^+^ T cell immunosurveillance in non-lymphoid tissues is dominated by resident populations. Whether CD4^+^ T cells use the same strategies to survey peripheral tissues is less clear. Here, examining the turnover of CD4^+^ T cells in transplanted duodenum in humans, we demonstrate that the majority of CD4^+^ T cells were still donor-derived one year after transplantation. In contrast to memory CD4^+^ T cells in peripheral blood, intestinal CD4^+^ T_RM_ cells expressed CD69 and CD161, but only a minor fraction expressed CD103. Functionally, intestinal CD4^+^ T_RM_ cells were very potent cytokine producers; the vast majority being polyfunctional T_H_1 cells, whereas a minor fraction produced IL-17. Interestingly, a fraction of intestinal CD4^+^ T cells produced granzyme-B and perforin after activation. Together, we show that the intestinal CD4^+^ T-cell compartment is dominated by resident populations that survive for more than 1 year. This finding is of high relevance for the development of oral vaccines and therapies for diseases in the gut.

## Introduction

Studies of mouse models of infection have shown that CD8^+^ T cells remain in peripheral tissues long after pathogen clearance (Masopust et al., 2001). These long-lived CD8^+^ T cells have limited potential to recirculate and have been termed resident memory T (T_RM_) cells (Masopust and Soerens, 2019; Mueller and Mackay, 2016; Szabo et al., 2019). Moreover, CD8^+^ T_RM_ cells show an extraordinary ability to mount rapid and potent *in situ* responses after infectious re-exposure (Beura et al., 2018; Park et al., 2018; Schenkel et al., 2013). The currently most established markers to identify CD8^+^ T_RM_ cells in barrier tissues are CD69 and CD103 (Bartolome-Casado et al., 2019; Mackay et al., 2013; Snyder et al., 2019). CD69 is rapidly upregulated after arrival into the tissue (Klonowski et al., 2004), and plays a key role preventing tissue egress by antagonizing sphingosine 1-phosphate receptor (S1PR1) (Skon et al., 2013). CD103 (also known as αE integrin) is highly expressed on intraepithelial lymphocytes (IELs) and the heterodimer αEβ7 binds E-cadherin on the surface of epithelial cells (Cepek et al., 1994; Schon et al., 1999), promoting the accumulation of IELs in the epithelium.

Although CD4^+^ T cells are more abundant than CD8^+^ T cells in most peripheral tissues (Sathaliyawala et al., 2013), studies to understand T_RM_ cell biology have mainly focused on CD8^+^ T cells. Over the last decade, CD4^+^ T_RM_ cells have been identified in lungs (Hondowicz et al., 2016; Teijaro et al., 2011), skin (Glennie et al., 2015; Watanabe et al., 2015) and the reproductive tract (Iijima and Iwasaki, 2014). However, the CD4^+^ T_RM_ population seems to be more heterogeneous and functionally plastic compared to CD8^+^ T_RM_ cells (Becattini et al., 2015; Brucklacher-Waldert et al., 2014), and whether CD4^+^ T cells in peripheral tissues are truly resident, non-circulatory cells is still a matter of debate (Carbone and Gebhardt, 2019).

CD103 as well as CD69 are induced by TGF-β, which is constitutively produced by gut epithelial cells (Zhang and Bevan, 2013). At human mucosal sites, most CD4^+^ T cells express CD69, but few express CD103 compared to CD8^+^ T_RM_ cells (Sathaliyawala et al., 2013), and both CD103^-^ and CD103^+^ CD4^+^ T subsets have been described in different tissues, such as lung (Oja et al., 2018; Snyder et al., 2019) and skin (Watanabe et al., 2015). Intestinal CD4^+^ T_RM_ cells have shown to play a critical role in protection against different pathogens, including *C. rodentium* (Bishu et al., 2019) and *Listeria* (Romagnoli et al., 2017) in mouse models. Although our knowledge about the role of intestinal CD4^+^ T-cell effector subsets in the pathogenesis of inflammatory bowel disease (IBD) (Kleinschek et al., 2009; Lamb et al., 2017; Zundler et al., 2019) and coeliac disease (Christophersen et al., 2019; Risnes et al., 2018) have substantially progressed over the last decade, our current understanding of CD4^+^ T-cell immunosurveillance and long-term persistence in the human intestine remains incomplete.

We have recently reported that the majority of CD8^+^ T cells persists for years in human small intestine (Bartolome-Casado et al., 2019), however, it is still unknown whether CD4^+^ T cells share these features with their CD8^+^ counterparts. Here, we present a comprehensive study of the longevity and phenotype of intestinal CD4^+^ T cells in humans. In a unique transplantation setting we followed the persistence of donor-derived CD4^+^ T cells in grafted duodenum over time and found that the majority of donor CD4^+^ T cells are maintained for at least one year in the graft. Furthermore, both CD103^-^ and CD103^+^ CD4^+^ T cell populations presented very similar turnover rates, suggesting that both constitute T_RM_ populations. Finally, we showed that the vast majority of both CD103^-^ and CD103^+^ CD4^+^ T_RM_ cells were polyfunctional T_H_1 cells and a fraction produced cytotoxic granules after activation.

## Results

### Human intestinal CD4^+^ T cells are phenotypically distinct from their circulating counterparts

To identify CD4^+^ T cells with a T_RM_-phenotype in the human small intestine (SI) we first studied the CD4^+^ T-cell compartment under steady state conditions. For this purpose we collected SI specimens from proximal duodenum-jejunum resections of patients undergoing pancreatic cancer surgery (Whipple procedure, n = 35; mean age 63yr; 16 female), and from donors and recipients during pancreatic-duodenal Tx (baseline samples, donors: n = 52; mean age 31yr; 24 female; patients: n = 36; mean age 41yr; 14 female). All tissue samples were evaluated by experienced pathologists and only histologically normal SI was included. Single-cell suspensions from epithelium and enzyme-digested lamina propria (LP) were obtained and analyzed by flow cytometry together with peripheral blood mononuclear cells (PBMCs) from the patients. To characterize the phenotypic profile of SI CD4^+^ T cells, we performed flow-cytometry analysis using a panel of antibodies that we recently implemented to study SI CD8^+^ T_RM_ cells (Bartolome-Casado et al., 2019). CD4^+^ T cells comprised almost 60% of LP T cells (with a CD4^+^: CD8^+^ ratio similar to PB), but constituted only 10% of T cells in the epithelium (**Figure S1A-B**). The relative distribution of T cell subsets was conserved in mucosal biopsies sampled up to 35 cm apart from the same intestinal resection (**Figure S1C**).

Applying a dimensionality reduction technique (UMAP, Uniform Manifold Approximation and Projection) on the compiled flow cytometry data we found that all SI CD4^+^ T cells clustered separate from PB CD4^+^ T cells (**Figure 1A, left**). The vast majority of the SI CD4^+^ T cells presented a CD45RA^-^ CD45RO^+^ L-Sel^-^ CCR7^-^ effector memory (T_EM_) phenotype (**Figure 1A-C**). In contrast, PB CD4^+^ T cells contained a substantial fraction of naïve (T_N_, CD45RO^-^ CD45RA^+^ CCR7^+^ L-Sel^+^) and central memory (T_CM_, CD45RO^+^ CD45RA^-^ CCR7^+^ L-Sel^+^) CD4^+^ T cells (**Figure 1A-C**). Virtually all SI CD4^+^ T cells expressed the T_RM_ marker CD69 whereas all PB CD4^+^ T cells were CD69-negative (**Figure 1A, C**). The SI CD4^+^ T cells were separated into three clusters based on their differential expression of CD103 and KLRG1 (**Figure 1A**). The population expressing CD103 comprised on average 18% of the LP and 66% of IE CD4^+^ T cells (**Figure 1C, left**), whereas KLRG1 was expressed by 26% and 5% of LP and IE CD4^+^ T cells, respectively (**Figure 1D**). PB CD4^+^ T cells were completely negative for CD103, however a fraction (mean 19%) of PB CD4^+^ T_EM_ cells expressed KLRG1 (**Figure 1A and D**). PB CD4^+^ T_EM_ and SI CD4^+^ T cells showed similar expression of PD1, CD127 (IL-7 receptor-α) and NKG2D. In contrast, CD28 was significantly higher expressed on PB CD4^+^ T_EM_ cells, whereas CD161 was expressed at higher levels on SI CD4^+^ T cells (**Figure 1A and D**). In addition, the immunomodulatory receptor CD244 (2B4) was expressed higher on the IE subset. In line with other reports (Kumar et al., 2017), we also found that the negative regulator CD101 was highly expressed by the SI CD4^+^ T cells (**Figure 1S-D**). Given that one of the SI CD4^+^ T-cell clusters was enriched in cells expressing the T_RM_ marker CD103 (**Figure 1A**), we examined the differential phenotypic profile of CD103^+^ and CD103^-^ CD4^+^ T cells in LP and in the epithelium. CD103^-^ CD4^+^ T cells presented a higher fraction of KLRG1 positive cells in both compartments, while IE CD103^+^ CD4^+^ T cells exhibited significantly higher expression of 2B4. Otherwise we found only small differences between the CD103^+^ and CD103^-^ subsets (representative histograms in **Figure 1E** and compiled data in **Figure S1E)**.

**Figure 1.**
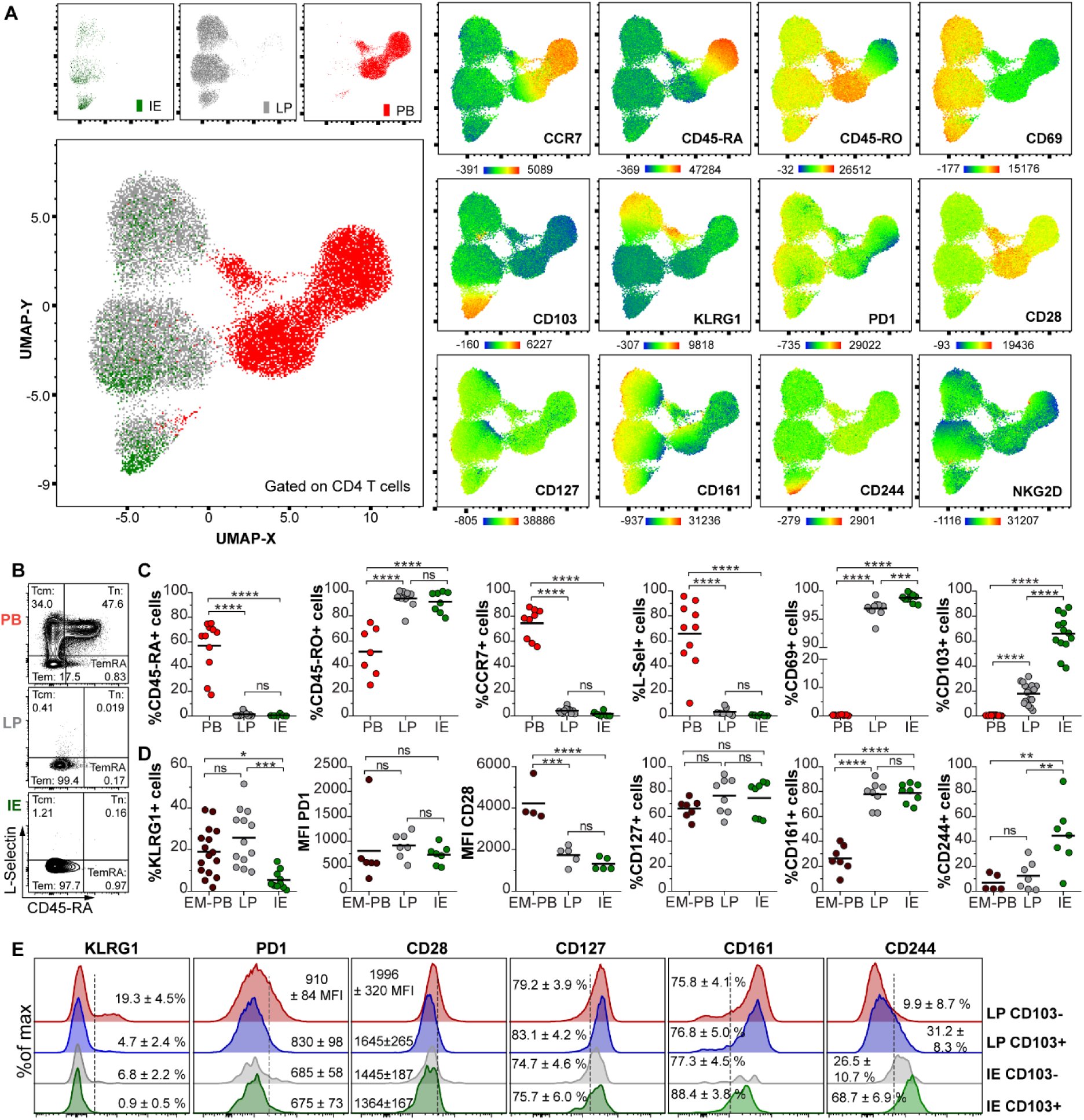
Human intestinal CD4^+^ T cells are phenotypically distinct from their circulating counterparts. **(A)** UMAP visualization after concatenation of flow-cytometric data from PB (red), LP (gray), and IE (green) CD4^+^ T cells, as described in (Bartolome-Casado et al., 2019). Representative of three samples. Map of the clusters and representation of each tissue compartment (left). Overlay of the UMAP clusters and the expression levels for each marker, color-coded based on the median fluorescence intensity values (MFI) (right). **(B)** Representative contour plots showing L-selectin and CD45RA expression on PB, LP and IE CD4^+^ T cells and classification of these cells into Tcm, central memory; Tem, effector memory; TemRA, effector memory re-expressing CD45RA; Tn, naïve. **(C)** Phenotypic comparison of total PB CD4^+^ T cells or **(D)** effector memory (EM) PB CD4^+^ T cells with intestinal LP, and IE CD4^+^ T cells. Compiled data for each marker are given and black bars indicate mean values. One-way ANOVA with Tukey’s multiple comparisons test. ns, not significant; *, P ≤ 0.05; **, P ≤ 0.01 ***, P ≤ 0.001; ****, P ≤ 0.0001. **(E)**. Representative histograms showing the differential phenotypic profile of intestinal CD103^−^ and CD103^+^ CD4^+^ T cells from LP and IE for several T_RM_-related markers. Mean values and SEM is provided. Compiled data of all the experiments are shown in **Figure S1C**.

Taken together, these results show that SI CD4^+^ T cells were clearly different from their blood counterparts, being CD69^+^ CD103^+^/^-^ CD161^+^ CD28^low^. The phenotype of IE CD4^+^ T cells was very similar to IE CD8^+^ cells; the majority being CD103^+^ KLRG1^-^ 2B4^+^.

### CD4^+^ T_RM_ cells persist for >1 yr in the transplanted SI

To directly examine the longevity of CD4^+^ T_RM_ cells in human SI, we assessed the long-term persistence of donor CD4^+^ T cells in endoscopic biopsies obtained from grafted duodenum at 3, 6 and 52 weeks after pancreatic-duodenal transplantation (Tx) of type I diabetic patients (Horneland et al., 2015). Only patients without histological or clinical signs of rejection were included (n=32). Most donors and recipients expressed different human leukocyte antigen (HLA) type I molecules rendering it possible to distinguish donor cells from incoming recipient cells in the graft by flow cytometry **(Figure 2A-B**). The CD103^-^ and CD103^+^ CD4^+^ T cells were analyzed separately. At 3 and 6 weeks, LP and IE CD4^+^ T cells exhibited very low replacement (median >85% donor cells), with no significant differences between the CD103^-^ and CD103^+^ CD4^+^ T subsets **(Figure 2B**). Importantly, also 1-yr after Tx the majority of SI CD4^+^ T cells in both the LP and IE compartments were donor-derived, the fraction being slightly higher for LP CD103^+^ compared to CD103^-^ CD4^+^ T cells (medians 77% and 60%, respectively). However, the majority of CD103^-^ CD4^+^ T cells were still of donor origin at 1 yr post-Tx, demonstrating that CD103 expression is not required for the persistence of CD4^+^ T_RM_ cells in human SI. In line with this, the turnover of both IE and LP CD103^+^ and CD103^-^ cells was highly correlated at 1 yr post-Tx (**Figure 2C**). Moreover, at 1 yr post-Tx the CD103^-^ and CD103^+^ CD4^+^ T-cell subsets contained a similar (or higher) proportion of donor cells compared to donor CD8^+^ T cell subsets (**Figure 2D**) (Bartolome-Casado et al., 2019). To confirm the persistence of donor CD4^+^ T cells we performed immunostaining with anti-CD3 and anti-CD4 antibodies combined with fluorescent *in situ* hybridization probes specific for X/Y-chromosomes on tissue sections where recipients and donors were of different gender and consistently observed donor-derived CD4^+^ T cells in the graft 1-yr after Tx (**Figure 2E**).

**Figure 2.**
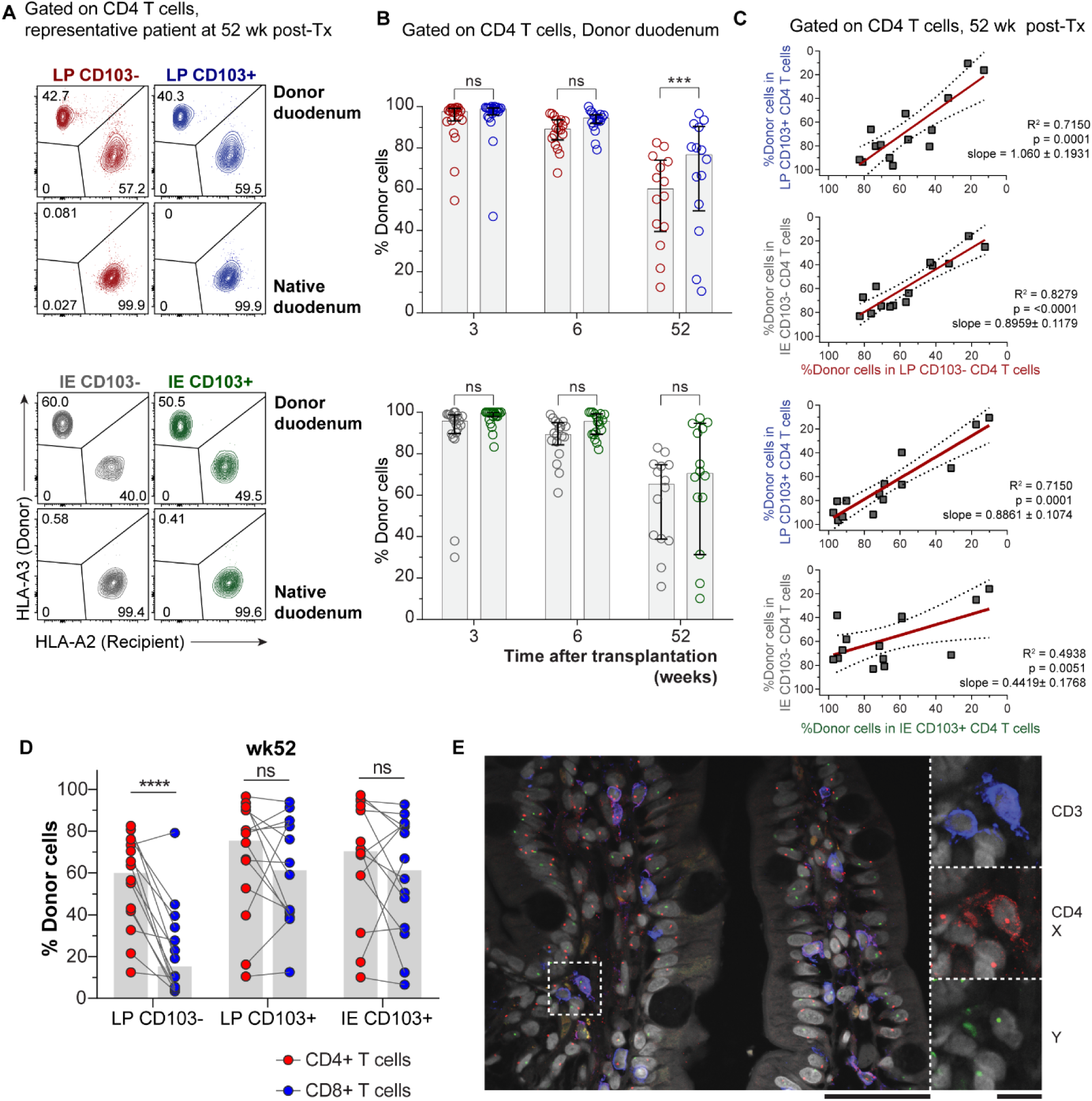
CD4^+^ T_RM_ cells persist for >1 yr in the transplanted SI. **(A)** Representative contour plots at 52 wk after Tx and **(B)** compile data for the fractions of donor-derived CD4^+^ T cells in LP CD103^−^ (red) and CD103^+^ (blue) compartments as well as IE CD103^−^ (grey) and CD103^+^ (green) at 3 (n = 20), 6 (n = 18), and 52 wk after Tx (n = 14) determined by HLA class I expression (as in A). Gray columns indicate median values. Statistical analysis was performed using two-way ANOVA for repeated measures (RM) with Tukey’s multiple comparisons test. ns, not significant; ***, P ≤ 0.001; **(C)** Pearson correlation of the percentages of donor-derived cells in the above mentioned LP and IE CD4^+^ T cell subsets 1-yr after Tx. Statistics performed using two-tailed P value (95% confidence interval, n = 14). **(D)** Frequencies of persisting donor cells in different subsets of CD4^+^ and CD8^+^ T cells from the same biopsies of donor duodenum at 52 weeks (wk) post-Tx. Data for CD8^+^ T cells has been previously published (Bartolome-Casado et al., 2019). Grey bars indicate median values. RM two-way ANOVA. ****, P ≤ 0.0001; ns, non-significant. **(E)** Representative confocal image of biopsies obtained from donor duodenum (male) of a female patient at one year post-transplantation. Tissue sections were stained with X/Y chromosome fluorescent in situ hybridization probes (Y, green; X, red) and antibodies against CD4 (red) and CD3 (blue). Hoechst (gray) stains individual nuclei. Scale bars, 50µm and 10µm, respectively.

These results showed that the majority of donor-derived SI CD4^+^ T cells persisted at least 1 yr (possibly years) in the tissue. However, to exclude effects of the surgical trauma, immunosuppressive treatment and leukocyte chimerism on the SI CD4^+^ T-cell population, we examined the absolute T-cell counts in SI over time. Serial tissue sections were stained for CD3 and CD8, scanned and counted. The density of CD4^+^ T cells was determined by subtracting the number of CD8^+^ cells from the total CD3^+^ cell count. We found that the overall density of both CD4^+^ and CD8^+^ T cells in Tx duodenum was stable throughout the 1-yr follow-up period (**Figure S2A-B**). Intracellular staining of single cell suspensions from Tx biopsies with the proliferation marker Ki67 showed few Ki67-positive cells among the donor CD4^+^ T cells (**Figure S2C-D**). The percentage of Ki67^+^ CD4^+^ T cells was similar to that seen in the native duodenum in Tx patients and in steady state controls (**Figure S2D**), indicating that proliferation did not contribute substantially to the large number of persisting donor CD4^+^ T cells in transplanted SI. Finally, we confirmed that the CD4^+^ T cells in the native (recipient) duodenum were exclusively recipient-derived (**Figure S3**), demonstrating that migration of donor cells out of the graft was not occurring.

In conclusion, these results show that the CD4^+^ T_RM_ cell population includes both CD103^-^ and CD103^+^ cells, and that CD4^+^ T_RM_ cells are at least as persistent as CD103^+^ CD8^+^ T_RM_ cells (Bartolome-Casado et al., 2019) in the transplanted SI.

### Incoming recipient CD4^+^ T cells undergo gradual phenotypic changes over time in transplanted duodenum

Transplanted SI gives us a unique opportunity to study the differentiation of recruited incoming CD4^+^ T cells and whether they acquire a T_RM_ phenotype in SI mucosa. To this end, we compared the expression of T_RM_ associated markers on donor- and recipient-derived LP CD4^+^ T cells from biopsies of transplanted duodenum over time. Already at 3 wk post-Tx, virtually all recipient LP CD4^+^ T cells expressed CD69 (**Figure 3A**). More than half of recipient CD4^+^ T cells expressed CD161 at 6 weeks and that was further increased at 1-yr post Tx to similar levels as donor CD4^+^ T cells (**Figure 3B**). CD103 was expressed on a minor subset of recipient-derived CD4^+^ T cells at both 6 and 52 weeks; slightly lower than that on donor CD4^+^ T cells (**Figure 3C, E**). In contrast, the fraction of KLRG1-positive cells within donor and recipient-derived CD4^+^ T cells remained almost unchanged (**Figure 3D-E)**. Similarly to the steady state conditions (**Figure 1A**), the majority of the LP CD4^+^ T cells were CD103^-^ KLRG1^-^ at all the time points regardless of their origin (**Figure 3E**). Furthermore, the turnover of donor LP CD103^-^ KLRG1^-^ and CD103^-^ KLRG1^+^ CD4^+^ T cells were very similar, evidenced by the high correlation of donor-derived cells within both subsets over time (**Figure 3F**).

**Figure 3.**
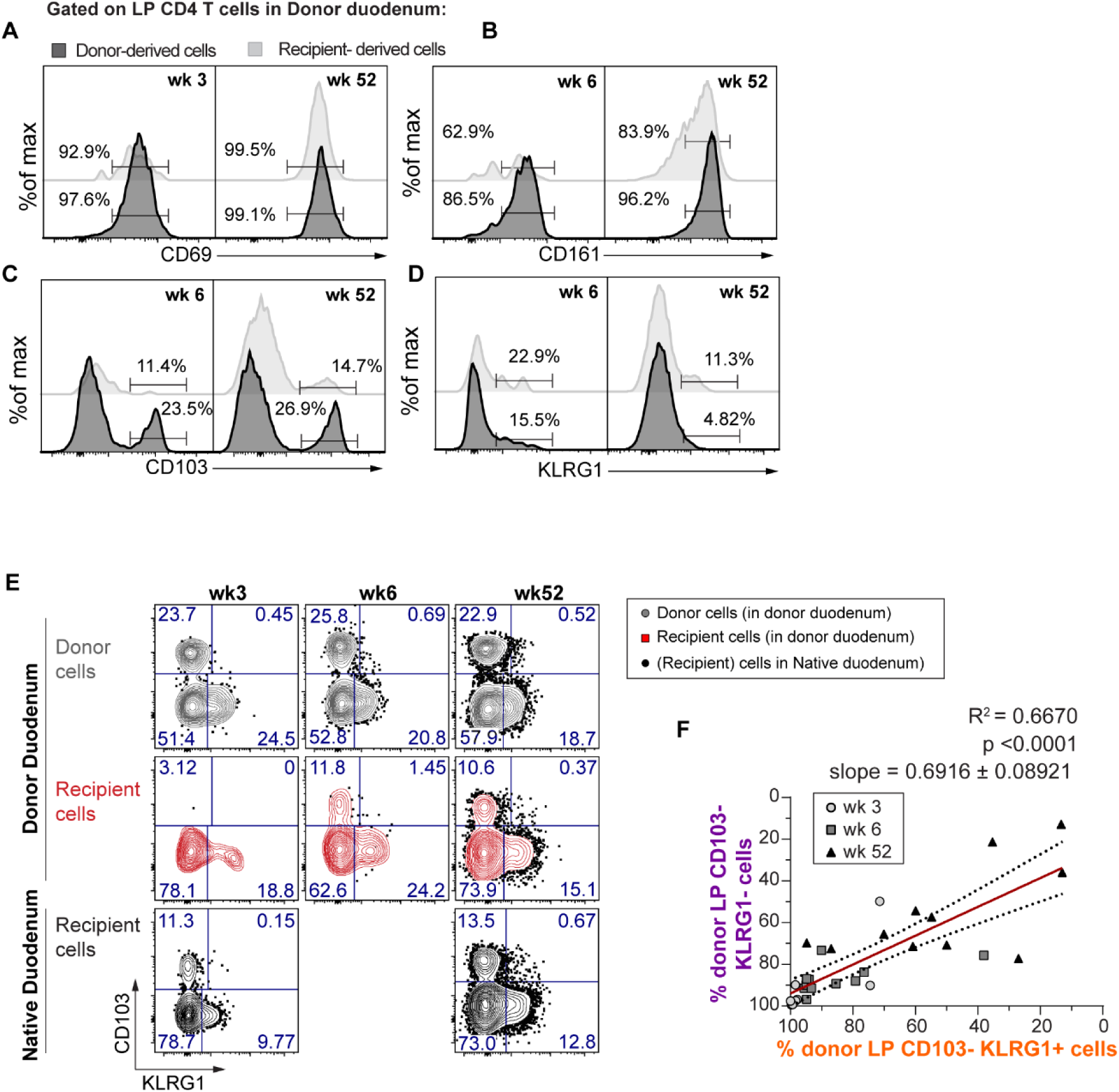
Incoming recipient CD4^+^ T cells undergo gradual phenotypic changes over time in transplanted duodenum. **(A-D)** Representative flow-cytometric analysis for the expression of different T_RM_ associated markers on donor-derived (black) and recipient-derived (light grey) LP CD4^+^ T cells from donor duodenum at the indicated weeks (wk) post-Tx. **(E)** Distribution of donor (grey) and recipient-derived (red) LP CD4^+^ T cells isolated from donor duodenum (above) and native duodenum (below, black) according to the expression of CD103 and KLRG1 at the indicated time-points after-Tx. **(F)** Pearson correlation of the percentages of donor-derived cells in LP CD103^-^ KLRG1^+^ and KLRG1^-^ CD4^+^ T cell subsets over time after Tx. Statistics performed using two-tailed P value (95% confidence interval, n = 32).

Together, we find that recipient CD4^+^ T cells recruited to the transplanted duodenum rapidly acquire phenotypic features similar to persistent donor CD4^+^ T cells (**Figure 2**), suggesting that they gradually differentiate into T_RM_ *in situ*.

### The majority of SI CD4^+^ T cells exhibits a polyfunctional T_H_1 profile

To examine the functional properties of SI CD4^+^ T cells we studied their cytokine expression profile and ability to produce cytotoxic granules. First, LP CD4^+^ T cells isolated from histologically normal SI were short-term stimulated with PMA and Ionomycin and intracellular staining was perform with antibodies targeting specific cytokines (**Table S1**). By flow-cytometric analysis we found that the majority of the LP CD4^+^ T cells, both CD103^-^ and CD103^+^, produced IFN-γ, IL-2 and TNF-α **(Figure 4A)**. Almost half of the cells produced all these three cytokines simultaneously **(Figure 4B-C)**, and we did not find significant differences between CD103^-^ KLRG1^+^ and KLRG1^-^ cells (**Figure S4A**). In contrast, triple-producing cells constituted only 4% of the memory CD4^+^ T cells in PB **(Figure 4B-C)**. Comparing the LP CD103^-^ and CD103^+^ subsets, we found significantly higher fraction of IL-17 and MIP1-β-producing cells within the CD103^+^ subset compared to CD103^-^ CD4^+^ T cell subset **(Figure 4A)**. Furthermore, CD103^+^ CD4^+^ T cells contained a higher fraction of IFN-γ^+^ IL-17^+^ double producing cells **(Figure 4D)**. In contrast, CD103^-^ CD4^+^ T cells presented higher numbers of IL-13-producing cells than their CD103^+^ counterparts, whereas comparable expression of IL-10 and IL-22 was found in the two subsets **(Figure 4A)**.

**Figure 4.**
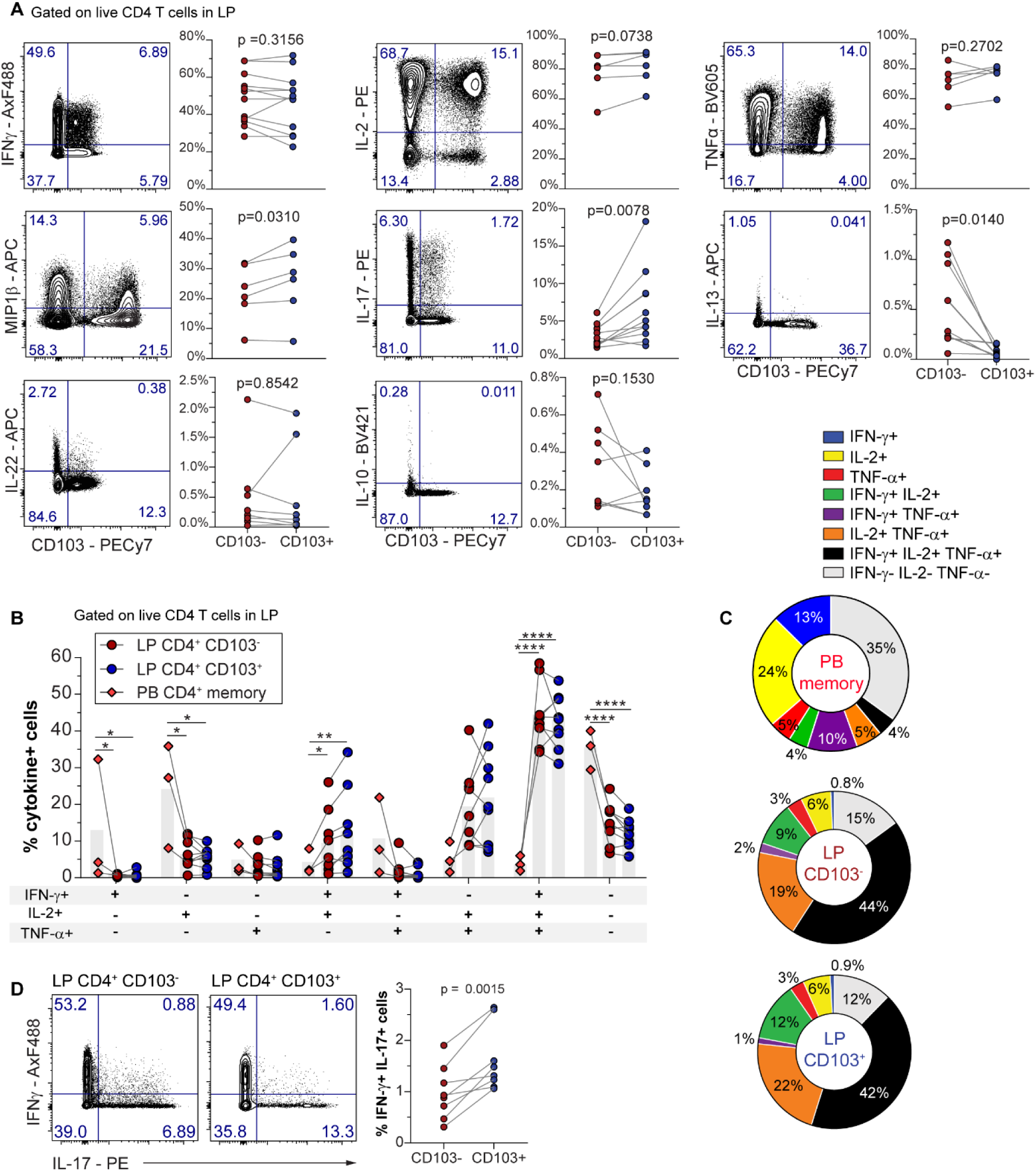
The majority of SI CD4^+^ T cells exhibits a polyfunctional T_H_1 profile. **(A)** Representative contour plots (left) and compiled data (right) showing PMA/ionomycin induced cytokine production by CD103^-^ compared to CD103^+^ LP CD4^+^ T cells. P values of paired t-test are displayed. **(B)** IFN-γ,IL-2 and TNF-α production by PB T_EM_ CD4^+^ T cells and intestinal LP CD103^−^, LP CD103^+^ CD4^+^ T cells. The bars indicate mean values. Statistics performed using one-way ANOVA for each combination of cytokines. **(C)** Relative fractions of each cytokine combination indicated in **(B)** by PB T_EM_ CD4^+^ T cells (n=3), and intestinal LP CD103^−^ (n=6) and LP CD103^+^ (n=6). CD4^+^ T cells represented on pie charts with color codes. Mean values of indicated experiments. **(D)** Representative contour plots (left) and compiled data (right) showing simultaneous IFN-γ and IL17 expression by LP CD103^-^ and CD103^+^ CD4^+^ T cells. Paired t-test.

Murine CD4^+^ T_RM_ cells have exhibited upregulation of granzyme-B upon reactivation with their cognate antigen (Beura et al., 2019). We therefore analyzed the capacity of SI CD4^+^ T cells to produce granzyme-B or perforin at the steady state and after stimulation with anti-CD3/CD28 beads. In the absence of stimulation, very few cells expressed these cytolytic proteins, however, both LP CD103^-^ and CD103^+^ subsets increased their expression of granzyme-B and perforin after activation **(Figure 5)**. We found a significantly higher proportion of granzyme-B producing cells within the LP CD103^+^ subset as compared to the CD103^-^ CD4^+^ T cell subset **(Figure 5)**. On the other hand, no significant differences were found in the activation-induced production of perforin between either subsets. **(Figure 5).** Comparing the KLRG1^+^ and KLRG1^-^ cells in the LP CD103^-^ compartment, we found higher basal levels of granzyme-B among the KLRG1^+^ cells (**Figure S4B**), but similar levels of granzyme-B and perforin after stimulation (**Figure S4B-C**).

**Figure 5.**
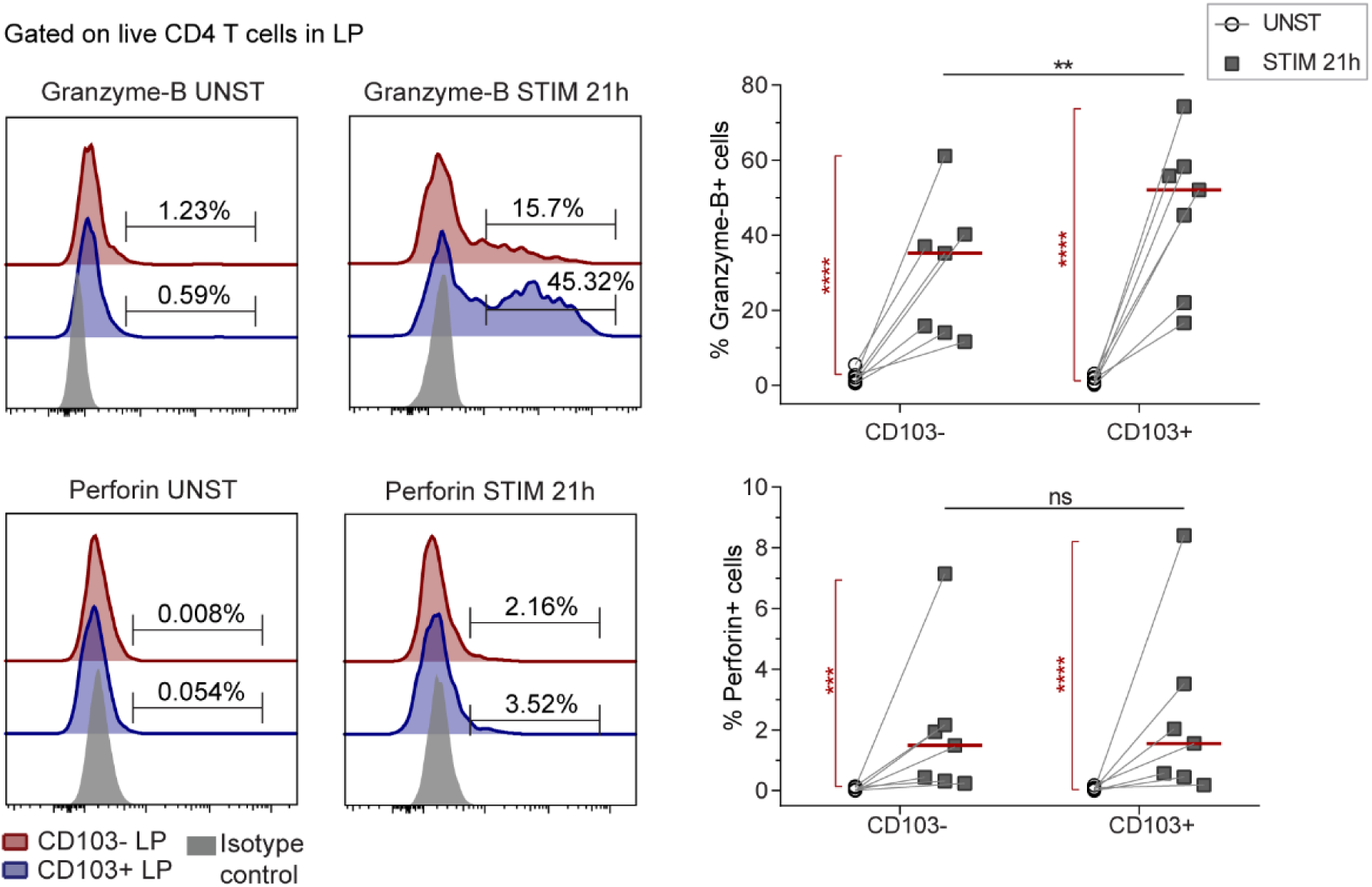
LP CD4^+^ T cells produce cytotoxic granules after stimulation. Representative flow-cytometric histogram (left) and compiled data (right) for the intracellular expression of granzyme-B and perforin in LP CD103^-^ and CD103^+^ CD4^+^ T cell subsets without (UNST) and after (STIM) stimulation with anti-CD3/CD28 beads for 21 h. Red lines indicate median values. Student’s t test was applied to compare the expression of cytotoxic mediators by both subsets (black horizontal lines), and by unstimulated *versus* 21h stimulated cells (red vertical lines and asterisks). **, P ≤ 0.01; ns, not-significant.

These data show that the majority of the SI CD4^+^ T_RM_ cells are polyfunctional T_H_1 cells, with a large fraction co-producing IFN-γ, IL-2 and TNF-α. A fraction of CD4^+^ T_RM_ cells also produces the cytotoxic proteins granzyme-B and perforin after stimulation.

## Discussion

Over the last years it has been demonstrated that immunosurveillance by memory CD8^+^ T cell in barrier tissues is largely mediated by durable, resident cell populations. However, whether memory CD4^+^ T cells use similar surveillance strategies is less clear (Carbone and Gebhardt, 2019; Homann et al., 2001; Snyder et al., 2019; Watanabe et al., 2015). Here, we show that the majority of CD4^+^ T cells are persistent for at least for 1 yr in the human SI mucosa, where they exhibit robust effector functions including polyfunctional T_H_1 responses.

There is conflicting evidence with regards to the long-term residency of memory CD4^+^ T cells in barrier tissues. Studies of CD4^+^ T_RM_ cells using parabiotic mice have suggested that CD4^+^ T-cell surveillance in the skin was dependent on continuous recirculation rather than permanent residency (Collins et al., 2016; Gebhardt et al., 2011). However, evidence of CD4^+^ T_RM_ cells persistence has been reported in other peripheral tissues, such as the reproductive mucosa and lung (Iijima and Iwasaki, 2014; Teijaro et al., 2011). Similarly, Beura et al. recently demonstrated that residency is the dominant mechanism of memory CD4^+^ T-cell immunosurveillance in non-lymphoid tissues, but they did not evaluate the longevity (Beura et al., 2019). Moreover, in a recent study Klicznik and colleagues discovered a population of skin CD103^+^ CD69^+^ CD4^+^ T cells that were able to downregulate CD69 expression and enter the circulation, indicating that some CD4^+^ T_RM_ cells may retain migratory potential (Klicznik et al., 2019).

In mouse models of infection, the number of antigen-specific memory CD4^+^ T cells in lymphoid and non-lymphoid tissues seem to decline faster than CD8^+^ T cells (Cauley et al., 2002; Homann et al., 2001), suggesting that memory CD4^+^ T cells are less durable. In line with these results, donor CD4^+^ T cells in lung transplanted patients were more rapidly lost than CD8^+^ T cells (Snyder et al., 2019). Here, we found that donor CD4^+^ T cells were maintained in duodenal grafts at equal or even higher numbers than CD103^+^ CD8^+^ T cells 1 yr after transplantation, without any change in cell density. In fact, in several patients more than 80% of the CD4^+^ T cells were donor-derived at 1 yr. It is reasonable to assume that the host response to an organ transplantation would increase, rather than decrease, the replacement kinetics of immune cells in the graft (Eguiluz-Gracia et al., 2016; Snyder et al., 2019; Zuber et al., 2016), together indicating that most CD4^+^ T cells in human SI are non-circulating, resident cells that most likely perpetuate for years.

Similar to intestinal CD8^+^ T_RM_ cells (Bartolome-Casado et al., 2019), we found that virtually all the SI CD4^+^ T cells expressed CD69 and CD161. However, unlike CD8^+^ T cells, only a minor fraction of LP CD4^+^ T cells expressed the αE integrin, CD103. While CD103^-^ CD8^+^ T cells very rapidly turned over in transplanted duodenum (**Figure 2E** and (Bartolome-Casado et al., 2019)), both CD103^-^ and CD103^+^ CD4^+^ T cells showed a similar persistence. These results show that retention of CD4^+^ T cells is independent of CD103, in line with previous reports (Romagnoli et al., 2017).

Like murine CD4^+^ T_RM_ cells (Iijima and Iwasaki, 2014; Romagnoli et al., 2017), the vast majority of LP CD4^+^ T_RM_ cells exhibited a polyfunctional T_H_1 profile, producing high amounts of IFN-γ, IL-2 and TNF-α. The fraction of polyfunctional T_H_1 cells among SI CD4^+^ T cells was much higher than among memory CD4^+^ T cells in blood. Furthermore, >40% of the CD4^+^ T_RM_ cells expressed granzyme-B after stimulation. These results show that SI CD4^+^ T_RM_ cells, like CD8^+^ T_RM_ cells, undergo tissue-specific changes that make them poised to provide robust T_H_1 immunity in response to reinfections (Beura et al., 2019; Pope et al., 2001). In addition to protection against pathogens (Romagnoli et al., 2017), long-lived CD4^+^ T cell responses to commensal bacteria have been found during acute gastrointestinal infection with *T. gondii* (Hand et al., 2012). Moreover, microbiota-specific CD4^+^ T cells have been identified in blood and intestinal biopsies from healthy humans (Hegazy et al., 2017), indicating that CD4^+^ T_RM_ cells may actively contribute to intestinal homeostasis through interactions with the microbiota.

We found that a fraction of CD4^+^ T_RM_ cells produced IL-17. T_H_17 cells play an important role in intestinal inflammatory disorders (Kleinschek et al., 2009; Ouyang et al., 2008; Yang et al., 2014; Zundler et al., 2019), however IL-17 is also critical for maintaining mucosal barrier integrity (Martinez-Lopez et al., 2019; Ouyang et al., 2008). Recently it was reported that, in contrast to inflammatory T_H_17 cells elicited by pathogens, gut commensal bacteria elicited tissue-resident homeostatic T_H_17 cells, which showed limited capacity to produce inflammatory cytokines (Omenetti et al., 2019). In our study only a very small percentage of T_H_17 cells co-produced the inflammatory cytokine IFN-γ, suggesting that the majority of SI T_H_17 cells during homeostasis are non-inflammatory cells that support barrier integrity. However, further studies are needed to understand the role of SI T_H_17 cells under homeostatic and inflammatory conditions.

Finally, we found, although marginally, that CD103^+^ T_RM_ cells contained higher fractions of IL-17 single- and IL-17/IFN-γ double-producing cells than their CD103^-^ counterparts. Moreover, CD103^-^ and CD103+ CD4^+^ T cells also showed subtle phenotypic differences regarding their expression of KLRG1, CD28 and 2B4. However, to what extent the CD103^+^ and CD103^-^ subsets represents distinct functional subsets needs further investigation.

In conclusion, we provide evidence that the majority of memory CD4^+^ T cells in the human SI are resident and may persist in the tissue for >1 year. This indicates that residency constitute the dominant mechanism for CD4^+^ memory T cell immunosurveillance in the human SI, and should be explored for the development of oral vaccines as well as for strategies to treat CD4^+^ T-cell mediated inflammatory intestinal diseases.

## Materials and Methods

### Patient samples

Small intestinal samples were either obtained during pancreatic cancer surgery (Whipple procedure, n = 35; mean age 63yr; range 40-81yr; 16 female), or from donors and/or patients during pancreas-duodenum transplantation (donors: n = 52; mean age 31yr; range 5-55yr; 24 female; patients: n = 36; mean age 41yr; range 25-60yr; 14 female) as described previously (Bartolome-Casado et al., 2019). Cancer patients receiving neoadjuvant chemotherapy were excluded from the study. Endoscopic biopsies from donor and patient duodenum were collected at 3, 6 and 52 weeks after transplantation. All tissue specimens were evaluated blindly by experienced pathologists, and only material with normal histology was included (Ruiz et al., 2010). All transplanted patients received a standard immunosuppressive regimen (Horneland et al., 2015), and patients showing clinical or histological signs of rejection or other complications, as well as patients presenting pre-transplant or *de novo* donor specific antibodies (DSA) were excluded from the study. Blood samples were collected at the time of the surgery and buffy coats from healthy donors (Oslo University Hospital). All participants gave their written informed consent. The study was approved by the Regional Committee for Medical Research Ethics in Southeast Norway and complies with the Declaration of Helsinki.

### Preparation of intestinal and peripheral blood single-cell suspensions

Intestinal resections were opened longitudinally and rinsed with PBS, and mucosa was dissected in strips off the submucosa. For microscopy, small mucosal pieces were fixed in 4% formalin and embedded in paraffin according to standard protocols. Intestinal mucosa was washed 3 times in PBS containing 2mM EDTA and 1% FCS at 37°C with shaking for 20 minutes and filtered through nylon 100-µm mesh to remove epithelial cells. Epithelial fractions in each washing step were pooled and filtered through 100-µm cell strainers (BD, Falcon). Epithelial cells in the EDTA fraction were depleted by incubation with anti-human epithelial antigen antibody (clone Ber-EP4, Dako) followed by anti-mouse IgG dynabeads (ThermoFisher) according to the manufacture’s protocol. De-epithelialized LP was minced and digested in complete RPMI medium (supplemented with 1% Pen/Strep) containing 0.25 mg/mL Liberase TL and 20 U/mL DNase I (both from Roche), stirring at 37°C for 1h. Digested tissue was filtered twice through 100-µm cell strainers and washed tree times in PBS. Purity of both IE and LP fractions was checked by flow-cytometry (Bartolome-Casado et al., 2019). Intestinal biopsies from transplanted patients were processed in the same way. PBMCs were isolated by Ficoll-based density gradient centrifugation (Lymphoprep™, Axis-Shield).

### Flow cytometry

Single cell suspensions of intestinal LP and IE fractions and PBMCs were stained using different multicolor combinations of directly conjugated monoclonal antibodies (**Table S1**). To assess the expression of L-Selectin on digested tissue, cells were rested for 12h at 37°C before the immunostaining. Replacement of donor cells in duodenal biopsies of HLA mismatched transplanted patients was assessed using different HLA type I allotype-specific antibodies targeting donor- and/or recipient-derived cells, and stroma cells were used as a control of specific staining. Dead cells were excluded based on propidium iodide staining (Molecular Probes, Life Technologies). For analysis of cytokine production, LP and IE cell suspensions were stimulated for 4h with control complete medium (RPMI supplemented with 10% FCS, 1% Pen/Strep) or phorbol-12-myristate-13-acetate PMA (1.5 ng/mL) and ionomycin (1µg/mL; both from Sigma-Aldrich) in the presence of monensin (Golgi Stop, BD Biosciences) added after 1h of stimulation to allow intracellular accumulation of cytokines. Cells were stained using the BD Cytofix/Cytoperm kit (BD Biosciences) according to the manufacturer’s instructions, and stained with antibodies against several cytokines (**Table S1**). For detection of cytotoxic granules, LP and IE cells were activated for 21h with anti-CD3/CD28 beads (Dynabeads, ThermoFisher) or control complete medium. For detection of intranuclear Ki67 expression the FoxP3/transcription factor staining buffer set was used according to the manufacturer’s instructions. eFluor-450 or eFluor-780 fixable viability dyes (eBioscience) were used prior any intracellular/intranuclear staining procedure. All samples were acquired on LSR Fortessa flow cytometer (BD Biosciences), using FACSDiva software (BD Biosciences). Single stained controls were prepared for compensation (UltraComp eBeads™, eBioscience), and gates were adjusted by comparison with FMO controls or matched isotype controls. Flow cytometry data were analyzed using FlowJo 10.4.2 (Tree Star). For **Figure 1A**, the expression of 16 phenotypic markers was analyzed at the single cell-level and compared for CD4^+^ T cells in PB, LP and IE (n=3) using the merge and calculation functions of Infinicyt software (Cytognos), as described in detail elsewhere (Pedreira et al., 2013). The population within the CD4^+^ T-cell gate was down-sampled for each compartment and exported to a new file, and then concatenated and subjected to UAMP analysis using the plugin integrated in FlowJo 10.5.3 as in (Bartolome-Casado et al., 2019). All experiments were performed at the Flow Cytometry Core Facility, Oslo University Hospital.

### Microscopy

Analysis of chimerism was performed as described previously (Landsverk et al., 2017). Briefly, formalin-fixed 4-μm sections were washed sequentially in xylene, ethanol, and PBS. Heat-induced epitope retrieval was performed by boiling sections for 20min in Dako buffer. Sections were incubated with CEP X SpectrumOrange/Y SpectrumGreen DNA Probes (Abbott Molecular Inc.) for 12h at 37°C before immunostaining according to standard protocol with anti-CD3 (Polyclonal; Dako), anti-CD4 (clone 1F6, Leica Biosystems) and secondary antibodies targeting rabbit IgG or mouse IgG2b conjugated to Alexa Fluor 647 and 555, respectively. Laser scanning confocal microscopy was performed on an Olympus FV1000 (BX61WI) system. Image z stacks were acquired at 1-μm intervals and combined using the Z project max intensity function in Image J (National Institutes of Health), and all microscopy images were assembled in Photoshop and Illustrator CC (Adobe).

CD8 and CD3 immunoenzymatic staining was performed on formalin-fixed 4-μm sections, dewaxed in xylene and rehydrated in ethanol, and prepared with Vulcan Fast red kit (Biocare Medical) following standard protocols. In brief, heat-induced antigen retrieval was performed in Tris/EDTA pH9 buffer (EnVision FLEX Dako kit, K8010), followed by staining with primary antibody (CD8 clone 4B11, Novocastra or CD3, polyclonal, Dako), secondary anti-mouse AP-conjugated antibody and incubation with substrate (Fast red chromogen, Biocare Medical). Slides were counterstained with hematoxylin and excess of dye was removed by bluing in ammoniac water. Tissue sections were scanned using Pannoramic Midi slide scanner (3DHISTECH) and counts generated with QuPath software (Bankhead et al., 2017).

### Statistical analysis

Statistical analyses were performed in Prism 8 (GraphPad Software). To assess statistical significance among SI CD4^+^ T cell subsets, data were analyzed by one-way ANOVA (standard or repeated measures, RM-ANOVA) followed by Tukey’s multiple comparison tests. Replacement data and distribution of CD4^+^ T cell subsets at different time points were analyzed by two-way ANOVA matching across subsets followed by Tukey’s multiple comparison tests. Correlations between replacement kinetics of different CD4^+^ T cell subsets were calculated using Pearson correlation with two-tailed p-value (95% confidence interval). P-values of <0.05 were considered significant.

## Abbreviations

IE: intraepithelial
LP: lamina propria
RPMI: Roswell Park Memorial Institute medium
SI: small intestine
T_RM_: resident memory T cell
Tx: pancreatic-duodenal transplantation (of diabetes mellitus patients)

## Acknowledgments

We are grateful to the staff at the Endoscopy Unit and the surgical staff; Christian Naper, Institute of Immunology, for providing HLA typing; the Confocal Microscopy and Flow Cytometry Core Facilities; all at Oslo University Hospital, Rikshospitalet.

## Author contributions

R. Bartolomé-Casado, O.J.B. Landsverk, E.S. Bækkevold, and F.L. Jahnsen conceived the project. R. Bartolomé-Casado, O.J.B. Landsverk, and S.K. Chauhan processed samples, designed and performed experiments, and analyzed data. R. Bartolomé-Casado prepared figures. F. Sætre and K.Thorvaldsen Hagen assisted with experiments and data analysis. S. Yaqub and R. Horneland coordinated recruitment of patients and collection of biopsies. S. Yaqub, R. Horneland, O. Øyen, and E.M. Aandahl performed surgery and provided samples. L. Aabakken performed endoscopy and provided endoscopic biopsies. R. Bartolomé-Casado and F.L. Jahnsen wrote the manuscript. O.J.B. Landsverk, F. Sætre and E.S. Baekkevold contributed to writing the manuscript, E.S. Baekkevold, and F.L. Jahnsen supervised the study.

## Supplementary Materials

**Table 1:**
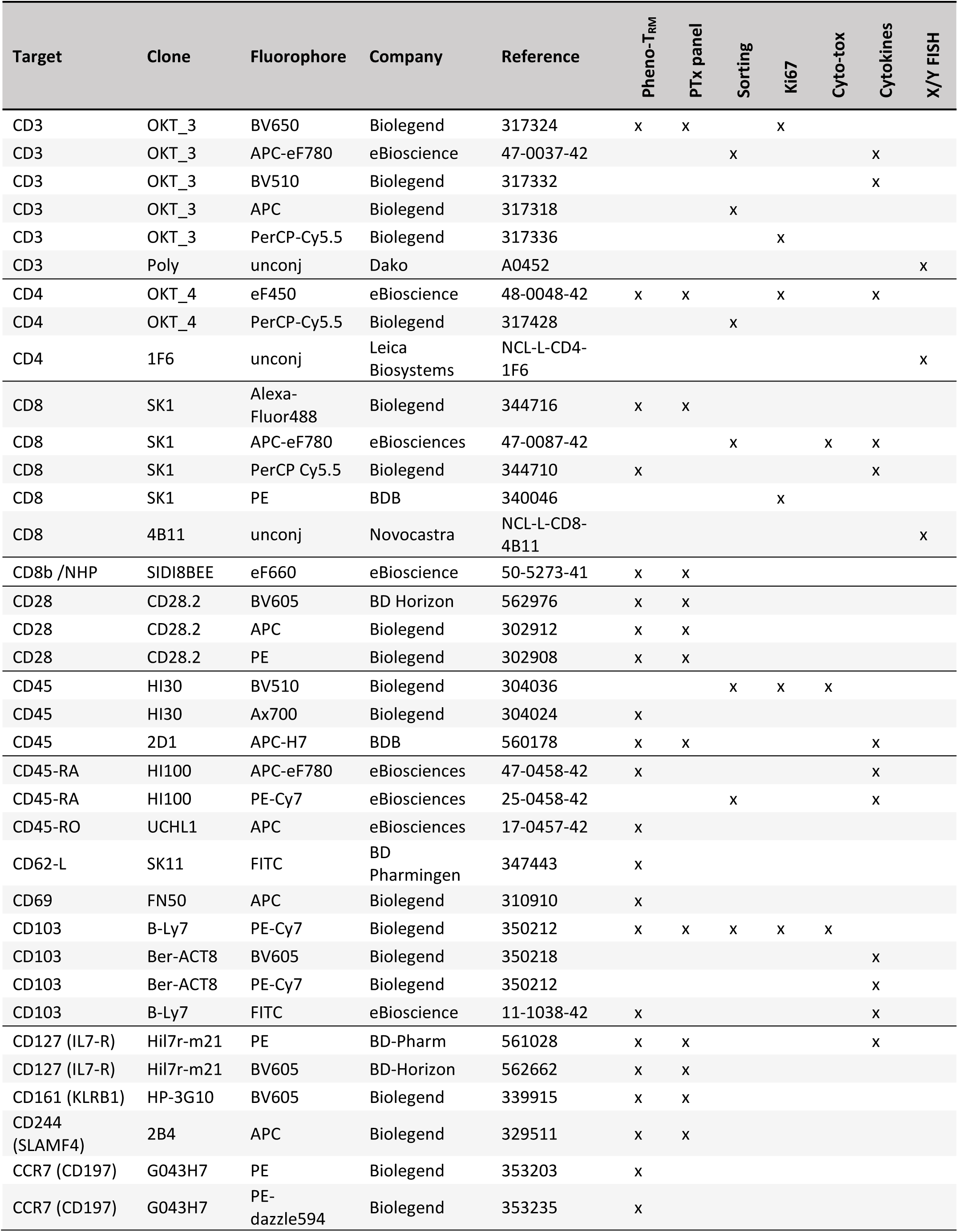
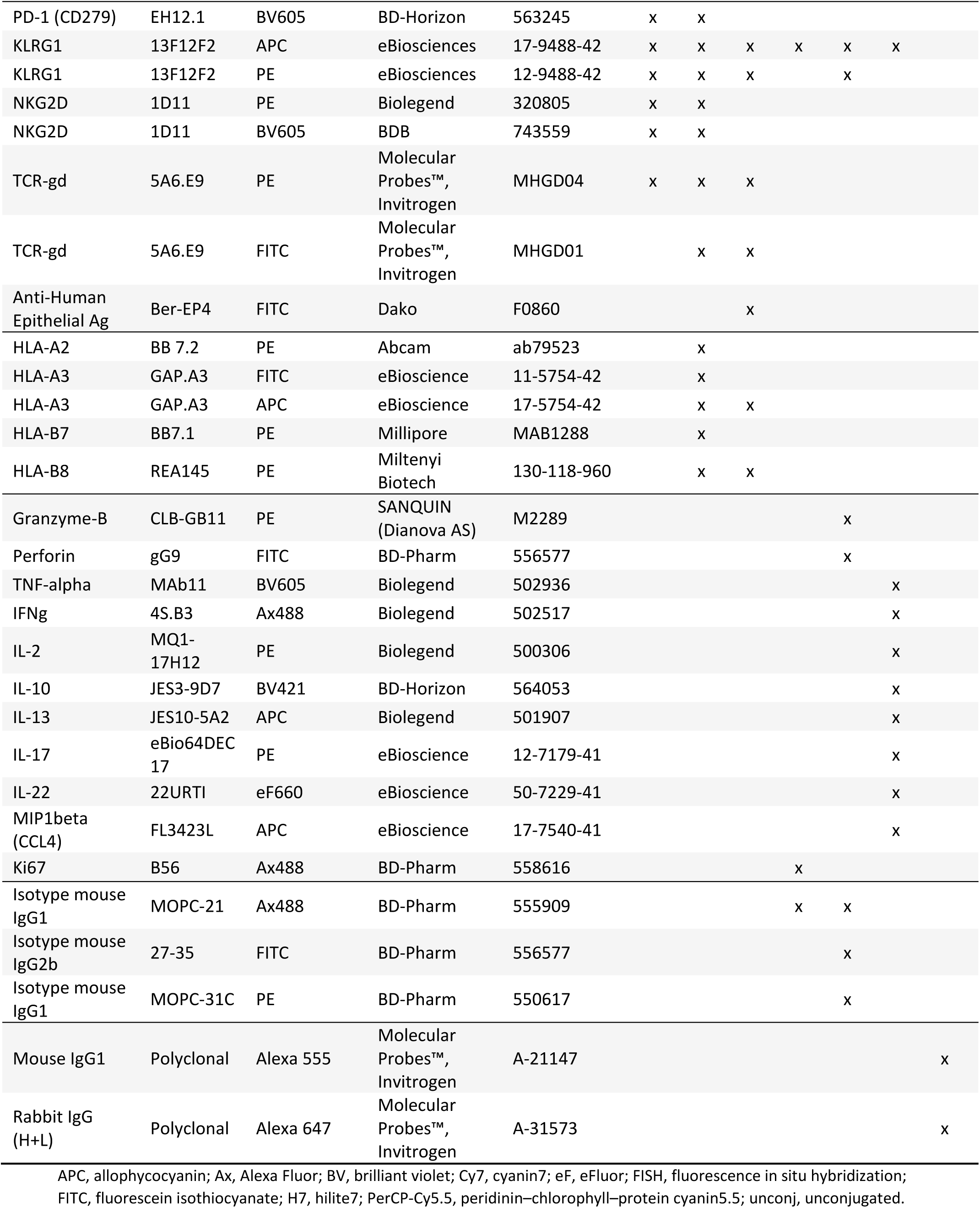
Antibodies used in the study.

**Figure S1.**
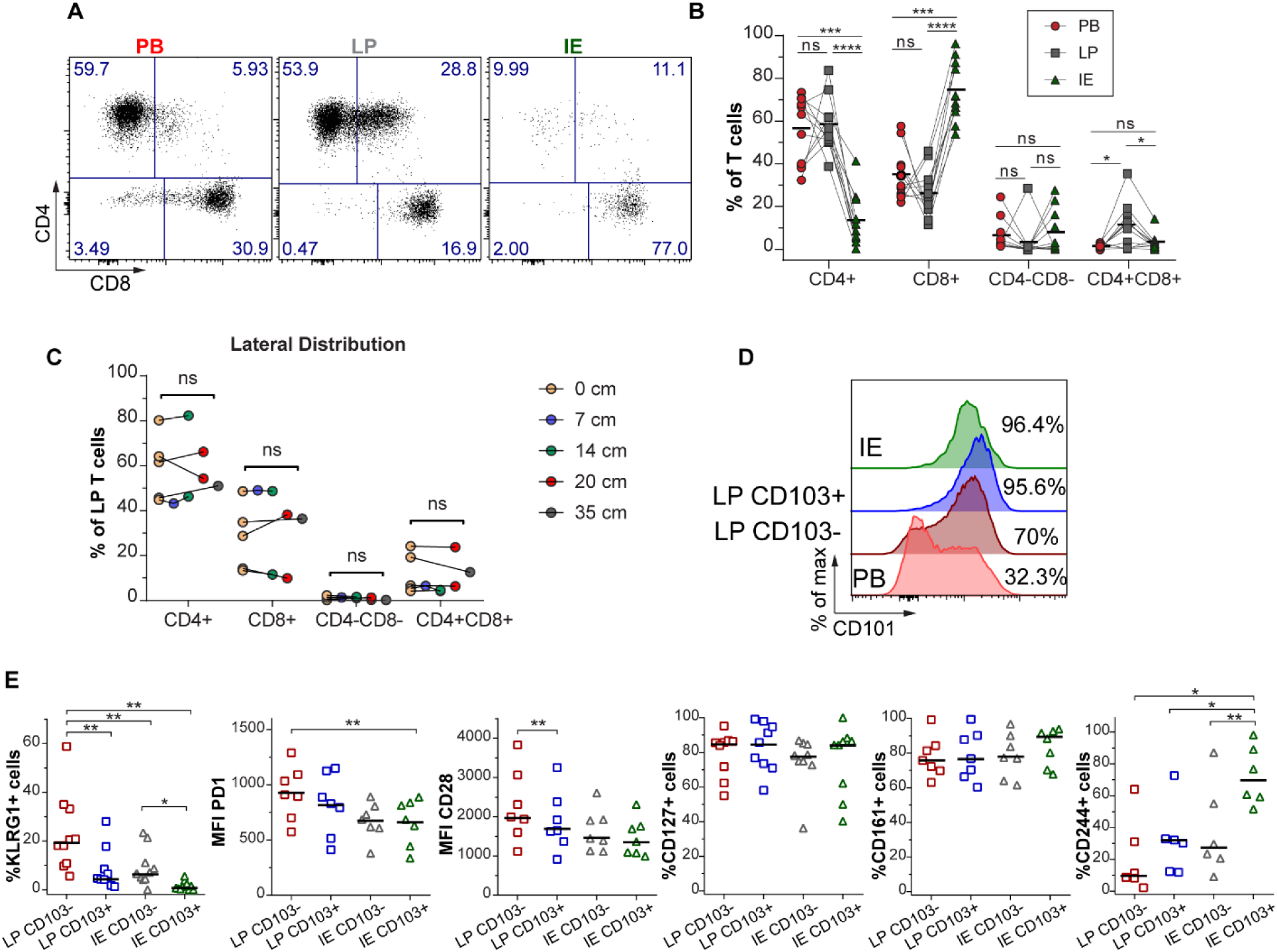
Extended phenotype of SI CD4^+^ T cells, accompanies Figure 1. **(A)** Representative flow-cytometric dot-plot and **(B)** aggregated data for the distribution of T cell subsets in PB, LP, and IE fractions collected from the same patient. Black lines indicate mean value. Statistics performed using two-way ANOVA, repeated measures matching both factors, and Tukey’s multiple comparison test. ns, not significant; *, P ≤ 0.05; ***, P ≤ 0.001; ****, P ≤ 0.0001. **(C)** Lengthwise representation of the CD4^+^ subsets in LP determined by flow cytometric analysis of biopsies taken at intervals along resected duodenum-proximal jejunum from individual subjects after Whipple procedure. n = 5; paired Student’s t test comparing 0 cm to the farthest distance. ns, not significant. **(D)** Representative histogram showing the expression of CD101 on PB, CD103^-^ and CD103^+^ LP and IE CD4^+^ T cells. **(E)** Percentage of positive cells or MFI values for various markers on intestinal-derived CD103^−^ and CD103^+^ CD4^+^ T cells from LP and IE. Black bars indicate median values. Statistical analysis was performed using repeated-measures one-way ANOVA with Tukey’s multiple comparisons test. *, P ≤ 0.05; **, P ≤ 0.01; all other comparisons are not significant.

**Figure S2.**
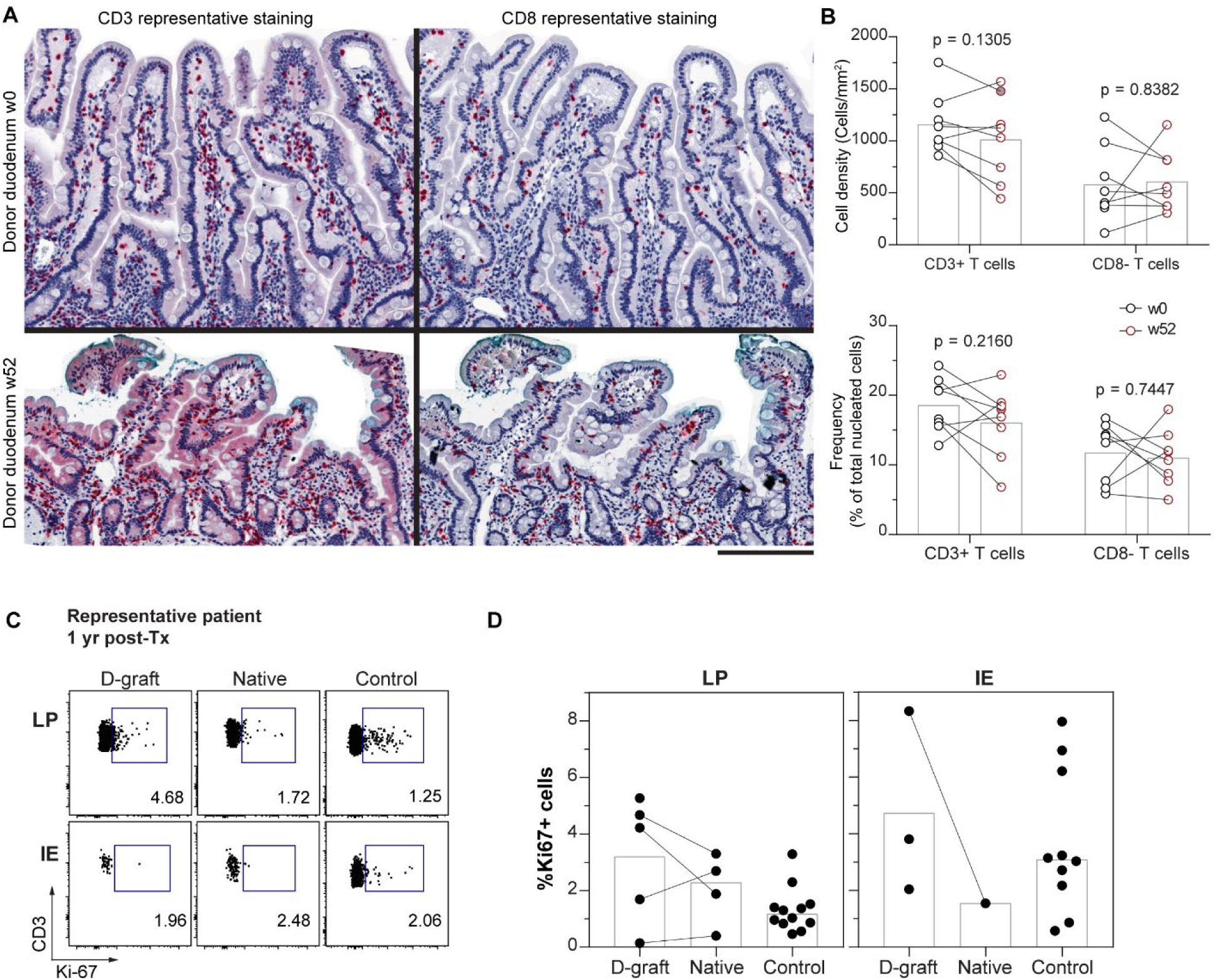
Turnover of CD4^+^ T cells in transplanted duodenum, accompanies Figure 2. **(A)** Representative immunohistochemistry staining of CD3^+^ cells (left) and CD8^+^ T cells (right) on tissue sections from donor duodenum at baseline (w0) and 1-yr after Tx (w52). Scale bar, 200µm. **(B)** Compile data of CD3^+^ and CD8^+^ T cell counts on tissue sections from donor duodenum of representative patients (n=8) at baseline (w0, black) and 1-yr after Tx (w52, red). Paired t-test. **(C)** Representative dot plot showing Ki67 expression in donor- or recipient-derived CD4^+^ T cells in LP and epithelium (IE) isolated from biopsies of donor or native duodenum 1-yr after Tx and control intestinal resections. **(D)** Compiled data for the percentages of Ki67-positive cells in donor- or recipient-derived CD4^+^ T cells in LP and IE at different time points after Tx in donor or native duodenum and in control intestinal resections.

**Figure S3.**
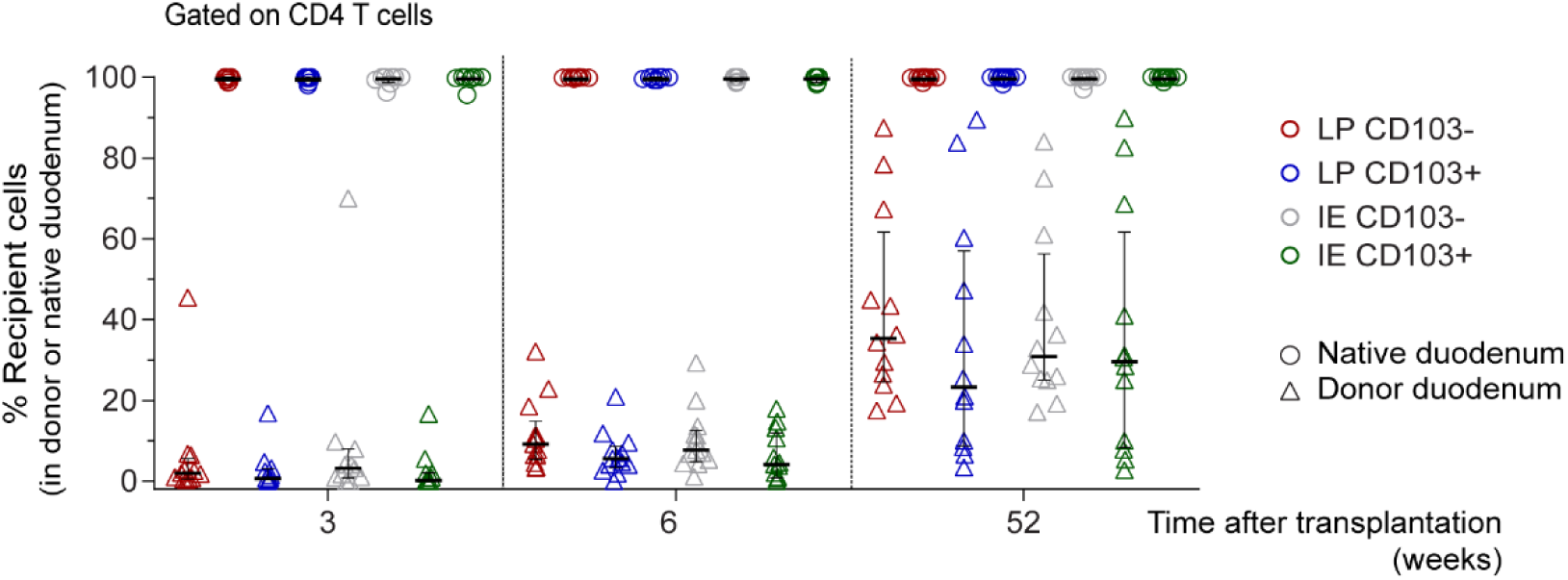
Absence of cross-contamination between donor and native (recipient) duodenum. Frequencies of recipient cells within each CD4^+^ T-cell subset in biopsies from donor and recipient (native) duodenum of the same patients at different time points after Tx. Black horizontal lines represent median values, and error bars show interquartile ranges.

**Figure S4.**
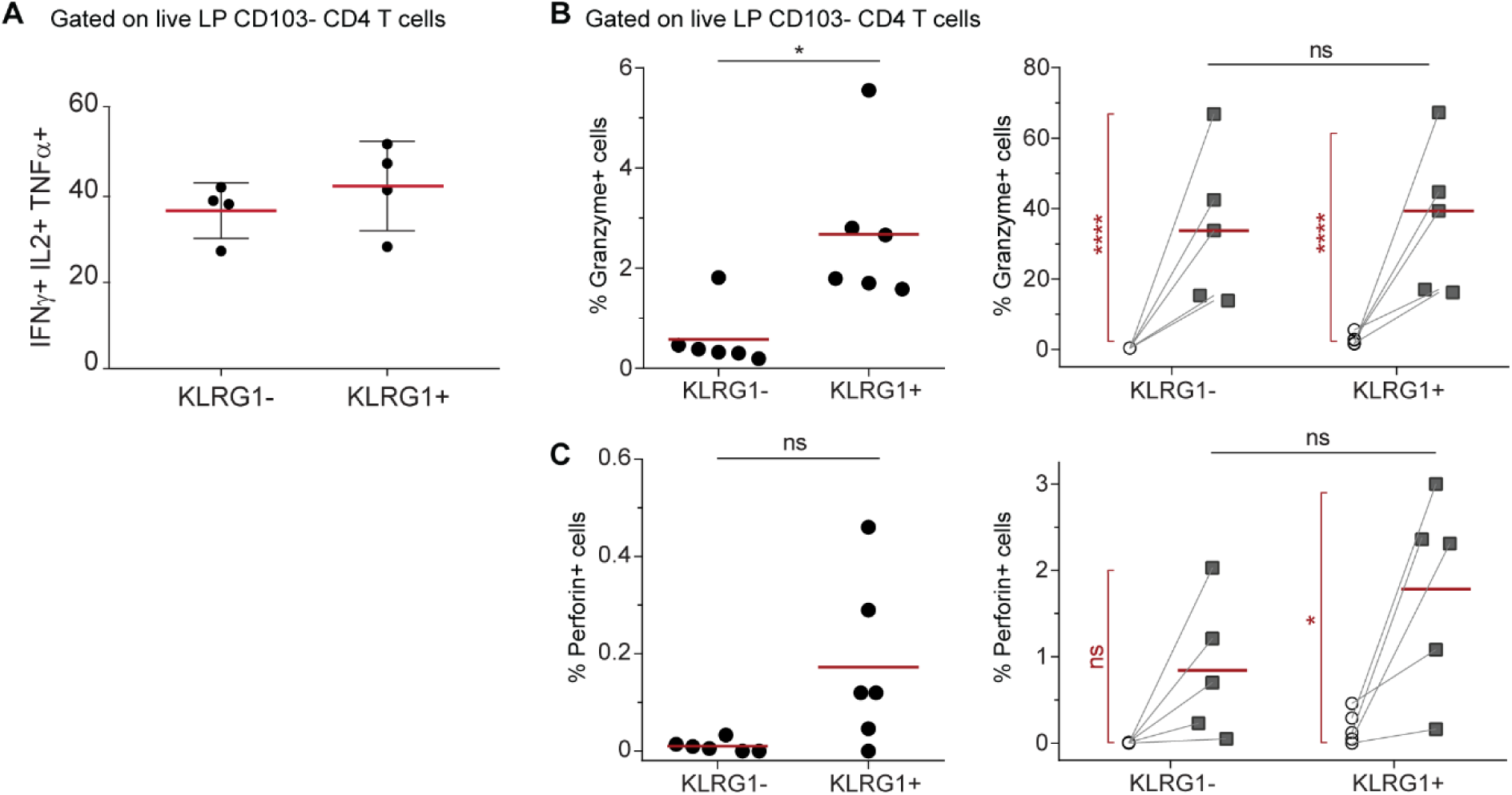
Comparative of the functional capabilities of LP CD103^-^ KLRG1^-^ and KLRG1^+^ CD4^+^ T cell subsets. **(A)** Frequency of polyfunctional (IL-2^+^, IFN-γ^+^ and TNF-α^+^) cells within LP CD103^-^ KLRG1^-^ and KLRG1^+^ subsets (n =4). **(B)** Frequencies of granzyme-B and **(C)** perforin positive cells in unstimulated samples (left) and after 21 h of activation with anti-CD3 beads (right) for the two subsets (KLRG1^+^, KLRG1^−^) of CD103^-^ CD4^+^ T cells in LP. Paired student’s t test between subsets (black) and unstimulated vs stimulated (red). Paired student’s t test was applied to compare the two subsets. ns, not significant; **, P ≤ 0.01.

## References

Bankhead, P., M.B. Loughrey, J.A. Fernandez, Y. Dombrowski, D.G. McArt, P.D. Dunne, S. McQuaid, R.T. Gray, L.J. Murray, H.G. Coleman, J.A. James, M. Salto-Tellez, and P.W. Hamilton. 2017. QuPath: Open source software for digital pathology image analysis. Sci Rep 7:16878.

Bartolome-Casado, R., O.J.B. Landsverk, S.K. Chauhan, L. Richter, D. Phung, V. Greiff, L.F. Risnes, Y. Yao, R.S. Neumann, S. Yaqub, O. Oyen, R. Horneland, E.M. Aandahl, V. Paulsen, L.M. Sollid, S.W. Qiao, E.S. Baekkevold, and F.L. Jahnsen. 2019. Resident memory CD8 T cells persist for years in human small intestine. J Exp Med

Becattini, S., D. Latorre, F. Mele, M. Foglierini, C. De Gregorio, A. Cassotta, B. Fernandez, S. Kelderman, T.N. Schumacher, D. Corti, A. Lanzavecchia, and F. Sallusto. 2015. T cell immunity. Functional heterogeneity of human memory CD4(+) T cell clones primed by pathogens or vaccines. Science 347:400–406.

Beura, L.K., N.J. Fares-Frederickson, E.M. Steinert, M.C. Scott, E.A. Thompson, K.A. Fraser, J.M. Schenkel, V. Vezys, and D. Masopust. 2019. CD4+ resident memory T cells dominate immunosurveillance and orchestrate local recall responses. The Journal of Experimental Medicine

Beura, L.K., J.S. Mitchell, E.A. Thompson, J.M. Schenkel, J. Mohammed, S. Wijeyesinghe, R. Fonseca, B.J. Burbach, H.D. Hickman, V. Vezys, B.T. Fife, and D. Masopust. 2018. Intravital mucosal imaging of CD8(+) resident memory T cells shows tissue-autonomous recall responses that amplify secondary memory. Nat Immunol 19:173–182.

Bishu, S., G. Hou, M. El Zaatari, S.R. Bishu, D. Popke, M. Zhang, H. Grasberger, W. Zou, R.W. Stidham, P.D.R. Higgins, J.R. Spence, N. Kamada, and J.Y. Kao. 2019. Citrobacter rodentium Induces Tissue-Resident Memory CD4(+) T Cells. Infect Immun 87:

Brucklacher-Waldert, V., E.J. Carr, M.A. Linterman, and M. Veldhoen. 2014. Cellular Plasticity of CD4+ T Cells in the Intestine. Front Immunol 5:488.

Carbone, F.R., and T. Gebhardt. 2019. Should I stay or should I go-Reconciling clashing perspectives on CD4(+) tissue-resident memory T cells. Sci Immunol 4:

Cauley, L.S., T. Cookenham, T.B. Miller, P.S. Adams, K.M. Vignali, D.A. Vignali, and D.L. Woodland. 2002. Cutting edge: virus-specific CD4+ memory T cells in nonlymphoid tissues express a highly activated phenotype. J Immunol 169:6655–6658.

Cepek, K.L., S.K. Shaw, C.M. Parker, G.J. Russell, J.S. Morrow, D.L. Rimm, and M.B. Brenner. 1994. Adhesion between epithelial cells and T lymphocytes mediated by E-cadherin and the alpha E beta 7 integrin. Nature 372:190–193.

Christophersen, A., E.G. Lund, O. Snir, E. Sola, C. Kanduri, S. Dahal-Koirala, S. Zuhlke, O. Molberg, P.J. Utz, M. Rohani-Pichavant, J.F. Simard, C.L. Dekker, K.E.A. Lundin, L.M. Sollid, and M.M. Davis. 2019. Distinct phenotype of CD4(+) T cells driving celiac disease identified in multiple autoimmune conditions. Nat Med 25:734–737.

Collins, N., X. Jiang, A. Zaid, B.L. Macleod, J. Li, C.O. Park, A. Haque, S. Bedoui, W.R. Heath, S.N. Mueller, T.S. Kupper, T. Gebhardt, and F.R. Carbone. 2016. Skin CD4(+) memory T cells exhibit combined cluster-mediated retention and equilibration with the circulation. Nat Commun 7:11514.

Eguiluz-Gracia, I., H.H. Schultz, L.I. Sikkeland, E. Danilova, A.M. Holm, C.J. Pronk, W.W. Agace, M. Iversen, C. Andersen, F.L. Jahnsen, and E.S. Baekkevold. 2016. Long-term persistence of human donor alveolar macrophages in lung transplant recipients. Thorax 71:1006–1011.

Gebhardt, T., P.G. Whitney, A. Zaid, L.K. Mackay, A.G. Brooks, W.R. Heath, F.R. Carbone, and S.N. Mueller. 2011. Different patterns of peripheral migration by memory CD4+ and CD8+ T cells. Nature 477:216–219.

Glennie, N.D., V.A. Yeramilli, D.P. Beiting, S.W. Volk, C.T. Weaver, and P. Scott. 2015. Skin-resident memory CD4+ T cells enhance protection against Leishmania major infection. J Exp Med 212:1405–1414.

Hand, T.W., L.M. Dos Santos, N. Bouladoux, M.J. Molloy, A.J. Pagan, M. Pepper, C.L. Maynard, C.O. Elson, 3rd, and Y. Belkaid. 2012. Acute gastrointestinal infection induces long-lived microbiota-specific T cell responses. Science 337:1553–1556.

Hegazy, A.N., N.R. West, M.J.T. Stubbington, E. Wendt, K.I.M. Suijker, A. Datsi, S. This, C. Danne, S. Campion, S.H. Duncan, B.M.J. Owens, H.H. Uhlig, A. McMichael, I.B.D.C.I. Oxford, A. Bergthaler, S.A. Teichmann, S. Keshav, and F. Powrie. 2017. Circulating and Tissue-Resident CD4(+) T Cells With Reactivity to Intestinal Microbiota Are Abundant in Healthy Individuals and Function Is Altered During Inflammation. Gastroenterology 153:1320–1337 e1316.

Homann, D., L. Teyton, and M.B. Oldstone. 2001. Differential regulation of antiviral T-cell immunity results in stable CD8+ but declining CD4+ T-cell memory. Nat Med 7:913–919.

Hondowicz, B.D., D. An, J.M. Schenkel, K.S. Kim, H.R. Steach, A.T. Krishnamurty, G.J. Keitany, E.N. Garza, K.A. Fraser, J.J. Moon, W.A. Altemeier, D. Masopust, and M. Pepper. 2016. Interleukin-2-Dependent Allergen-Specific Tissue-Resident Memory Cells Drive Asthma. Immunity 44:155–166.

Horneland, R., V. Paulsen, J.P. Lindahl, K. Grzyb, T.J. Eide, K. Lundin, L. Aabakken, T. Jenssen, E.M. Aandahl, A. Foss, and O. Oyen. 2015. Pancreas transplantation with enteroanastomosis to native duodenum poses technical challenges--but offers improved endoscopic access for scheduled biopsies and therapeutic interventions. Am J Transplant 15:242–250.

Iijima, N., and A. Iwasaki. 2014. T cell memory. A local macrophage chemokine network sustains protective tissue-resident memory CD4 T cells. Science 346:93–98.

Kleinschek, M.A., K. Boniface, S. Sadekova, J. Grein, E.E. Murphy, S.P. Turner, L. Raskin, B. Desai, W.A. Faubion, R. de Waal Malefyt, R.H. Pierce, T. McClanahan, and R.A. Kastelein. 2009. Circulating and gut-resident human Th17 cells express CD161 and promote intestinal inflammation. J Exp Med 206:525–534.

Klicznik, M.M., P.A. Morawski, B. Höllbacher, S.R. Varkhande, S.J. Motley, L. Kuri-Cervantes, E. Goodwin, M.D. Rosenblum, S.A. Long, G. Brachtl, T. Duhen, M.R. Betts, D.J. Campbell, and I.K. Gratz. 2019. Human CD4+CD103+ cutaneous resident memory T cells are found in the circulation of healthy individuals. Science Immunology 4:

Klonowski, K.D., K.J. Williams, A.L. Marzo, D.A. Blair, E.G. Lingenheld, and L. Lefrançois. 2004. Dynamics of Blood-Borne CD8 Memory T Cell Migration In Vivo. Immunity 20:551–562.

Kumar, B.V., W. Ma, M. Miron, T. Granot, R.S. Guyer, D.J. Carpenter, T. Senda, X. Sun, S.H. Ho, H. Lerner, A.L. Friedman, Y. Shen, and D.L. Farber. 2017. Human Tissue-Resident Memory T Cells Are Defined by Core Transcriptional and Functional Signatures in Lymphoid and Mucosal Sites. Cell Rep 20:2921–2934.

Lamb, C.A., J.C. Mansfield, G.W. Tew, D. Gibbons, A.K. Long, P. Irving, L. Diehl, J. Eastham-Anderson, M.B. Price, G. O’Boyle, D.E.J. Jones, S. O’Byrne, A. Hayday, M.E. Keir, J.G. Egen, and J.A. Kirby. 2017. alphaEbeta7 Integrin Identifies Subsets of Pro-Inflammatory Colonic CD4+ T Lymphocytes in Ulcerative Colitis. J Crohns Colitis 11:610–620.

Landsverk, O.J., O. Snir, R.B. Casado, L. Richter, J.E. Mold, P. Reu, R. Horneland, V. Paulsen, S. Yaqub, E.M. Aandahl, O.M. Oyen, H.S. Thorarensen, M. Salehpour, G. Possnert, J. Frisen, L.M. Sollid, E.S. Baekkevold, and F.L. Jahnsen. 2017. Antibody-secreting plasma cells persist for decades in human intestine. J Exp Med 214:309–317.

Mackay, L.K., A. Rahimpour, J.Z. Ma, N. Collins, A.T. Stock, M.L. Hafon, J. Vega-Ramos, P. Lauzurica, S.N. Mueller, T. Stefanovic, D.C. Tscharke, W.R. Heath, M. Inouye, F.R. Carbone, and T. Gebhardt. 2013. The developmental pathway for CD103(+)CD8+ tissue-resident memory T cells of skin. Nat Immunol 14:1294–1301.

Martinez-Lopez, M., S. Iborra, R. Conde-Garrosa, A. Mastrangelo, C. Danne, E.R. Mann, D.M. Reid, V. Gaboriau-Routhiau, M. Chaparro, M.P. Lorenzo, L. Minnerup, P. Saz-Leal, E. Slack, B. Kemp, J.P. Gisbert, A. Dzionek, M.J. Robinson, F.J. Ruperez, N. Cerf-Bensussan, G.D. Brown, D. Bernardo, S. LeibundGut-Landmann, and D. Sancho. 2019. Microbiota Sensing by Mincle-Syk Axis in Dendritic Cells Regulates Interleukin-17 and -22 Production and Promotes Intestinal Barrier Integrity. Immunity 50:446–461 e449.

Masopust, D., and A.G. Soerens. 2019. Tissue-Resident T Cells and Other Resident Leukocytes. Annu Rev Immunol

Masopust, D., V. Vezys, A.L. Marzo, and L. Lefrancois. 2001. Preferential localization of effector memory cells in nonlymphoid tissue. Science 291:2413–2417.

Mueller, S.N., and L.K. Mackay. 2016. Tissue-resident memory T cells: local specialists in immune defence. Nat Rev Immunol 16:79–89.

Oja, A.E., B. Piet, C. Helbig, R. Stark, D. van der Zwan, H. Blaauwgeers, E.B.M. Remmerswaal, D. Amsen, R.E. Jonkers, P.D. Moerland, M.A. Nolte, R.A.W. van Lier, and P. Hombrink. 2018. Trigger-happy resident memory CD4(+) T cells inhabit the human lungs. Mucosal Immunol 11:654–667.

Omenetti, S., C. Bussi, A. Metidji, A. Iseppon, S. Lee, M. Tolaini, Y. Li, G. Kelly, P. Chakravarty, S. Shoaie, M.G. Gutierrez, and B. Stockinger. 2019. The Intestine Harbors Functionally Distinct Homeostatic Tissue-Resident and Inflammatory Th17 Cells. Immunity

Ouyang, W., J.K. Kolls, and Y. Zheng. 2008. The biological functions of T helper 17 cell effector cytokines in inflammation. Immunity 28:454–467.

Park, S.L., A. Zaid, J.L. Hor, S.N. Christo, J.E. Prier, B. Davies, Y.O. Alexandre, J.L. Gregory, T.A. Russell, T. Gebhardt, F.R. Carbone, D.C. Tscharke, W.R. Heath, S.N. Mueller, and L.K. Mackay. 2018. Local proliferation maintains a stable pool of tissue-resident memory T cells after antiviral recall responses. Nat Immunol 19:183–191.

Pedreira, C.E., E.S. Costa, Q. Lecrevisse, J.J. van Dongen, A. Orfao, and C. EuroFlow. 2013. Overview of clinical flow cytometry data analysis: recent advances and future challenges. Trends Biotechnol 31:415–425.

Pope, C., S.K. Kim, A. Marzo, D. Masopust, K. Williams, J. Jiang, H. Shen, and L. Lefrancois. 2001. Organ-specific regulation of the CD8 T cell response to Listeria monocytogenes infection. J Immunol 166:3402–3409.

Risnes, L.F., A. Christophersen, S. Dahal-Koirala, R.S. Neumann, G.K. Sandve, V.K. Sarna, K.E. Lundin, S.W. Qiao, and L.M. Sollid. 2018. Disease-driving CD4+ T cell clonotypes persist for decades in celiac disease. J Clin Invest 128:2642–2650.

Romagnoli, P.A., H.H. Fu, Z. Qiu, C. Khairallah, Q.M. Pham, L. Puddington, K.M. Khanna, L. Lefrancois, and B.S. Sheridan. 2017. Differentiation of distinct long-lived memory CD4 T cells in intestinal tissues after oral Listeria monocytogenes infection. Mucosal Immunol 10:520–530.

Ruiz, P., H. Takahashi, V. Delacruz, E. Island, G. Selvaggi, S. Nishida, J. Moon, L. Smith, T. Asaoka, D. Levi, A. Tekin, and A.G. Tzakis. 2010. International grading scheme for acute cellular rejection in small-bowel transplantation: single-center experience. Transplant Proc 42:47–53.

Sathaliyawala, T., M. Kubota, N. Yudanin, D. Turner, P. Camp, J.J. Thome, K.L. Bickham, H. Lerner, M. Goldstein, M. Sykes, T. Kato, and D.L. Farber. 2013. Distribution and compartmentalization of human circulating and tissue-resident memory T cell subsets. Immunity 38:187–197.

Schenkel, J.M., K.A. Fraser, V. Vezys, and D. Masopust. 2013. Sensing and alarm function of resident memory CD8(+) T cells. Nat Immunol 14:509–513.

Schon, M.P., A. Arya, E.A. Murphy, C.M. Adams, U.G. Strauch, W.W. Agace, J. Marsal, J.P. Donohue, H. Her, D.R. Beier, S. Olson, L. Lefrancois, M.B. Brenner, M.J. Grusby, and C.M. Parker. 1999. Mucosal T lymphocyte numbers are selectively reduced in integrin alpha E (CD103)-deficient mice. J Immunol 162:6641–6649.

Skon, C.N., J.Y. Lee, K.G. Anderson, D. Masopust, K.A. Hogquist, and S.C. Jameson. 2013. Transcriptional downregulation of S1pr1 is required for the establishment of resident memory CD8(+) T cells. Nature Immunology 14:1285-+.

Snyder, M.E., M.O. Finlayson, T.J. Connors, P. Dogra, T. Senda, E. Bush, D. Carpenter, C. Marboe, L. Benvenuto, L. Shah, H. Robbins, J.L. Hook, M. Sykes, F. D’Ovidio, M. Bacchetta, J.R. Sonett, D.J. Lederer, S. Arcasoy, P.A. Sims, and D.L. Farber. 2019. Generation and persistence of human tissue-resident memory T cells in lung transplantation. Sci Immunol 4:

Szabo, P.A., M. Miron, and D.L. Farber. 2019. Location, location, location: Tissue resident memory T cells in mice and humans. Sci Immunol 4:

Teijaro, J.R., D. Turner, Q. Pham, E.J. Wherry, L. Lefrancois, and D.L. Farber. 2011. Cutting edge: Tissue-retentive lung memory CD4 T cells mediate optimal protection to respiratory virus infection. J Immunol 187:5510–5514.

Watanabe, R., A. Gehad, C. Yang, L.L. Scott, J.E. Teague, C. Schlapbach, C.P. Elco, V. Huang, T.R. Matos, T.S. Kupper, and R.A. Clark. 2015. Human skin is protected by four functionally and phenotypically discrete populations of resident and recirculating memory T cells. Sci Transl Med 7:279ra239.

Yang, J., M.S. Sundrud, J. Skepner, and T. Yamagata. 2014. Targeting Th17 cells in autoimmune diseases. Trends Pharmacol Sci 35:493–500.

Zhang, N., and M.J. Bevan. 2013. Transforming growth factor-beta signaling controls the formation and maintenance of gut-resident memory T cells by regulating migration and retention. Immunity 39:687–696.

Zuber, J., B. Shonts, S.P. Lau, A. Obradovic, J. Fu, S. Yang, M. Lambert, S. Coley, J. Weiner, J. Thome, S. DeWolf, D.L. Farber, Y. Shen, S. Caillat-Zucman, G. Bhagat, A. Griesemer, M. Martinez, T. Kato, and M. Sykes. 2016. Bidirectional intragraft alloreactivity drives the repopulation of human intestinal allografts and correlates with clinical outcome. Sci Immunol 1:

Zundler, S., E. Becker, M. Spocinska, M. Slawik, L. Parga-Vidal, R. Stark, M. Wiendl, R. Atreya, T. Rath, M. Leppkes, K. Hildner, R. Lopez-Posadas, S. Lukassen, A.B. Ekici, C. Neufert, I. Atreya, K. van Gisbergen, and M.F. Neurath. 2019. Hobit- and Blimp-1-driven CD4(+) tissue-resident memory T cells control chronic intestinal inflammation. Nat Immunol

